# A systemic circadian nicotinic acid riboside (NaR) signal engages the unfolded protein response and adipogenesis via the prefoldin complex

**DOI:** 10.64898/2026.03.04.709493

**Authors:** Ivan Vlassakev, Christina Savva, Liying Zhou, Danilo Ritz, Alexander Schmidt, Cholsoon Jang, Amir Ata Saei, Paul Petrus

## Abstract

Daily light–dark cycles impose predictable environmental fluctuations that require coordinated temporal regulation of cellular physiology. This coordination is mediated by the circadian clock, which operates as a network of tissue oscillators; however, the molecular signals that convey circadian information between organs remain incompletely defined. Here, we identify nicotinic acid riboside (NaR) as a circulating metabolite whose rhythmicity depends on the liver clock. In differentiating 3T3-L1 adipocytes, NaR engages unfolded protein response (UPR) gene programs and modulates adipogenic competence. Proteome-wide stability profiling implicates the prefoldin complex as a molecular target of NaR signaling, linking NaR exposure to altered proteostasis. Functionally, NaR-induced UPR signaling converges on the adipogenic transcription factor CEBPA, which is a central regulator of adipogenesis. Importantly, sustained NaR exposure suppresses adipocyte lipid deposition, whereas temporally restricted NaR stimulation enhances adipogenesis, indicating that NaR acts in a time-dependent manner. Together, these findings identify NaR as a liver clock–controlled circulating metabolite that couples systemic circadian metabolism to adipocyte proteostasis and differentiation, revealing a mechanism by which temporal metabolic signals shape tissue-specific physiological outcomes.

**Highlights:**

- The circadian clock is required for rhythmic regulation of circulating nicotinic acid riboside (NaR).
- The liver clock is sufficient to generate NaR rhythmicity.
- NaR engages the prefoldin complex to regulate unfolded protein response signaling and *Cebpa* expression.
- Time-dependent NaR exposure differentially regulates CEBPA levels and adipocyte lipid deposition.

## Introduction

Life on Earth evolved under predictable 24-hour light–dark cycles, driving the emergence of circadian mechanisms that temporally organize physiology and metabolism.^1,2^ In mammals, this organization is mediated by a genetically encoded molecular clock that aligns cellular processes with environmental time cues. At the molecular level, circadian rhythms arise from transcriptional–translational feedback loops in which the activating complex Brain and Muscle ARNT-Like 1 (BMAL1) together with Circadian Locomotor Output Cycles Kaput (CLOCK) or Neuronal PAS Domain Protein 2 (NPAS2) drives rhythmic gene expression, including that of its own repressors, the Period (PER) and Cryptochrome (CRY) proteins.^3^ These feedback loops generate near-24-hour oscillations in gene expression across virtually all tissues, providing each nucleated cell with an intrinsic time-keeping mechanism. While the molecular architecture of the circadian clock is well defined, increasing evidence indicates that circadian physiology emerges from coordinated interactions between tissue clocks rather than from isolated oscillators, and how circadian timing information is communicated between organs to ensure coherent systemic metabolic regulation remains incompletely understood.

Physiological processes such as systemic metabolism exhibit robust temporal coherence across tissues and can be dynamically reprogrammed by environmental challenges.^4,5^ This coherence is maintained by tissue clocks that form a coordinated inter-organ network aligning systemic physiology with external time cues.^6^ Recent studies have begun to reveal the complexity of this network, demonstrating that coordinated circadian regulation emerges from interactions between multiple tissue clocks rather than from isolated oscillators.^7–12^ These interactions are shaped by intrinsic clock function as well as behavioral, environmental, and physiological factors, underscoring the dynamic nature of systemic circadian regulation.^13,14^

We previously showed that the brain clock alone is sufficient to restore the majority of circulating metabolite rhythms, and that restricting feeding to the active phase similarly reinstates aspects of metabolic rhythmicity in clock-deficient mice ^7^. However, a subset of circulating metabolites fails to regain rhythmicity in the absence of peripheral tissue clocks, and the identity and physiological roles of these metabolites remain poorly understood. These clock-dependent metabolites are likely regulated through the integration of feeding cues by peripheral clocks^15^ and may function as systemic signals that coordinate energy metabolism across tissues.^10^ While multiple metabolic pathways contribute to circadian homeostasis^16^ accumulating evidence highlights a central role for tryptophan metabolism and the associated nicotinamide adenine dinucleotide (NAD^⁺^) pathway in linking metabolic state to circadian regulation.^17–19^

Here, we demonstrate that metabolites within the tryptophan and NAD^⁺^ pathways require an intact circadian clock system to maintain systemic oscillations. Among NAD^⁺^ intermediates, nicotinic acid riboside (NaR) exhibits robust rhythmicity in circulation, which is abolished in clock-deficient mice and accompanied by chronically elevated NaR levels. To define the cellular programs responsive to NaR, we performed transcriptomic profiling in differentiating adipocytes. This unbiased analysis revealed selective induction of unfolded protein response (UPR)–associated gene programs. Subsequent proteome-wide stability profiling implicated the prefoldin complex as a molecular target of NaR signaling. Functionally, NaR differentially regulates adipocyte lipid accumulation depending on temporal exposure, promoting lipid storage under pulsatile stimulation but not during continuous treatment. Together, these findings identify NaR as a liver clock–controlled circulating metabolite that links systemic circadian metabolism to adipocyte proteostasis and differentiation state.

## Results

### The role of the clock system in integrating feeding cues to control circulating metabolic rhythms

To identify circulating metabolic rhythms regulated by the molecular clock system, we reanalyzed our previously published circadian serum metabolomics dataset generated from night-fed wild-type (WT) and whole-body *Bmal1* knockout (KO) mice.^7^ Rhythmicity analyses were performed using the DryR framework to identify clock-dependent oscillations (**Figure 1A**). Of the 923 detected serum metabolites, 376 exhibited rhythmicity in both WT and KO mice (DryR model 4), whereas 86 metabolites were rhythmic exclusively in WT mice (DryR model 3), indicating clock-dependent control (**Figure 1B and figure S1A**). Analysis of metabolite class distributions revealed that feeding-driven rhythms largely mirrored the overall composition of detected metabolites (**Figure S1B**), consistent with a dominant influence of feeding–fasting cycles on systemic metabolic oscillations.^7,15^ In contrast, clock-dependent rhythmic metabolites displayed a distinct class distribution, characterized by enrichment of xenobiotics and cofactors/vitamins and a relative depletion of lipids (**Figure S1B**). Notably, within the cofactor/vitamin class, quinolinic acid and nicotinic acid riboside (NaR), two metabolites linked to tryptophan–NAD^⁺^ metabolism, emerged as clock-dependent candidates (**Figure 1C**)

**Figure 1.**
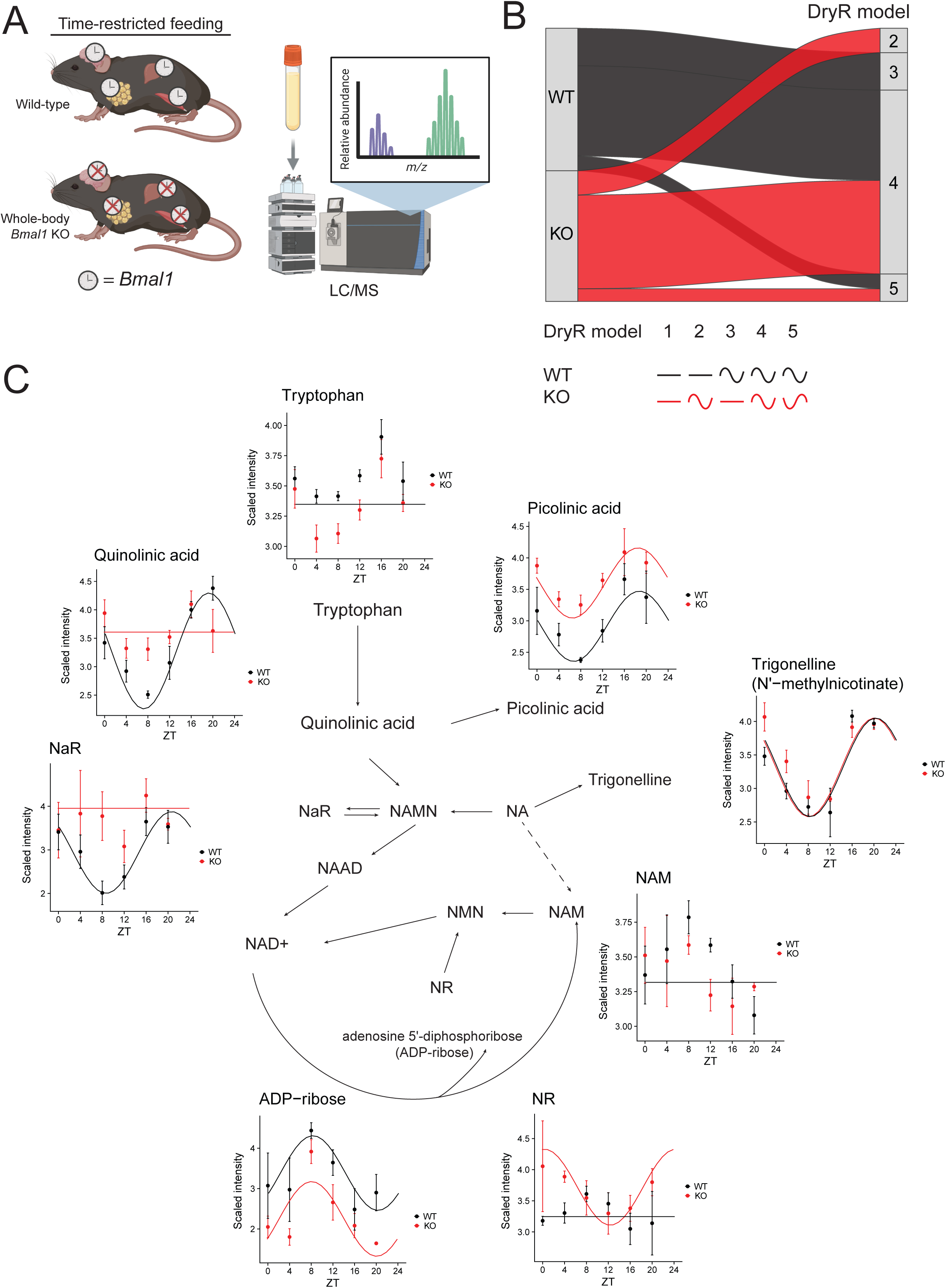
Circadian integration of feeding cues coordinates systemic metabolic rhythms. A) Schematic of experimental design (Created with BioRender.com). Four biological replicates of serum per genotype were sampled at each time point across the circadian cycle and analyzed using Liquid chromatography-mass spectrometry (LC/MS). B) Alluvial plot representing the post-hoc rhythmicity analyses performed using the DryR framework. C) Scaled intensity of serum detected tryptophan-NAD^+^ pathway-associated metabolites from WT (black) and whole-body *Bmal1*-KO (red) mice.

Our observations prompted us to focus on the tryptophan-NAD^⁺^ pathway. Examination of individual metabolites revealed distinct genotype-dependent rhythmic patterns. Quinolinic acid and NaR were the only metabolites detected in the serum that exhibited clock-dependent oscillations, whereas trigonelline, picolinic acid, and ADP-ribose remained rhythmic in both WT and KO mice, although the latter two displayed genotype-dependent mesor differences. In contrast, nicotinamide riboside (NR) was rhythmic exclusively in KO mice, while tryptophan and nicotinamide (NAM) showed no circadian regulation (**Figure 1C**). Together, these data indicate that only a specific subset of metabolites within the tryptophan and NAD^⁺^ pathways exhibit clock-dependent rhythmicity in circulation.

### The hepatic clock is sufficient to integrate feeding cues and restore liver NaR rhythmicity

The liver is a primary site for tryptophan and NAD^⁺^ metabolism.^20,21^ We therefore asked whether the hepatic clock is sufficient to integrate feeding cues and restore NaR rhythmicity. To address this, we reanalyzed our previously published liver metabolomics dataset comprising wild-type (WT), whole-body *Bmal1* knockout (KO), and hepatocyte-specific *Bmal1*-reconstituted (LRE) mice (**Figure 2A**).^15^ In LRE mice, circadian clock function is restored selectively in hepatocytes, while remaining absent in other tissues. All animals were maintained under time-restricted feeding conditions, with food access confined to the active phase (ZT12 to ZT0), enabling assessment of hepatic clock sufficiency in the context of controlled feeding–fasting cues.

**Figure 2.**
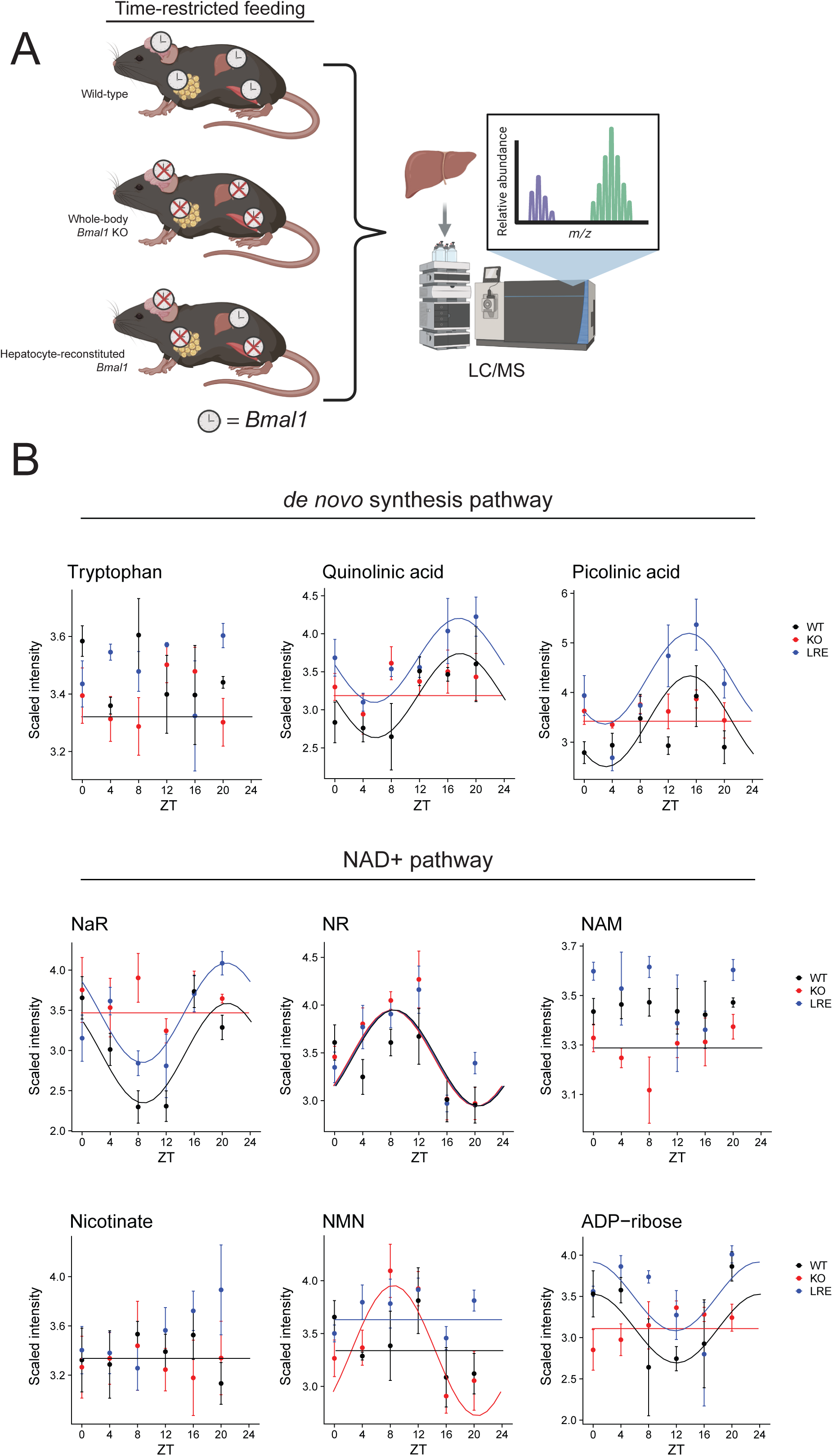
The liver clock alone integrates feeding cues to restore Nicotinic acid riboside (NaR) rhythms. A) Schematic of experimental design (Created with BioRender.com). Four biological replicates of serum per genotype were sampled at each time point across the circadian cycle and analyzed using Liquid chromatography-mass spectrometry (LC/MS). B) Scaled intensity of liver detected tryptophan-NAD+ pathway-associated metabolites from WT (black), whole-body Bmal1-KO (red) and hepatocyte-reconstituted *Bmal1* (LRE, blue) mice.

Across the 894 metabolites detected in the liver, approximately one third exhibited clock-independent rhythmicity (**Figure S2A and B**), underscoring the dominant influence of time restricted feeding on liver metabolism. In addition, 73 metabolites displayed liver clock–dependent oscillations (**Figure S2B**), indicating the role of the clock system in contributing to metabolic rhythms in a targeted manner. Upon hepatocyte-specific *Bmal1* reconstitution, rhythmicity was gained in relation to KO for a broad range of metabolites spanning amino acids, carbohydrates, nucleotides, and cofactors/vitamins, consistent with the liver clock functioning as a central metabolic regulator that integrates feeding-derived signals to control metabolic oscillations.^15^ In contrast, most metabolites that remained rhythmic exclusively in WT mice were lipids (**Figure S2B**), suggesting that oscillations in lipid metabolism require network dependent cross-tissue clock coordination.

We next examined the detected metabolites in serum within the tryptophan and NAD^⁺^ pathways. Among these, NaR, ADP-ribose, quinolinic acid, and picolinic acid regained rhythmicity upon hepatic *Bmal1* reconstitution (**Figure 2B**). In contrast, hepatic nicotinamide riboside (NR) oscillated independently of clock status, while nicotinamide mononucleotide (NMN) was rhythmic only in livers of KO mice (**Figure 2B**). Nicotinate, NAM, and tryptophan did not exhibit rhythmicity. Together, these data demonstrate that, the hepatic clock is sufficient to restore rhythmicity of both quinolinic acid and NaR, the two metabolites within the tryptophan-NAD^⁺^ pathway being clock-dependent in both serum and liver. While systemic regulation of quinolinic acid is linked to brain toxicity,^22^ NaR supplementation was recently shown to sustain systemic NAD^⁺^levels in aged mice.^20^ Thus, we focused our downstream investigations on the role of NaR.

### NaR regulates the unfolded protein response

We next asked whether the systemic NaR levels contribute to controlling biological functions in distant organs. We recently demonstrated that the liver clock plays a central role in modulating adipocyte rhythmicity.^12^ In addition, given the established importance of NAD^⁺^ metabolism for adipose tissue function,^23–26^ we chose to investigate the role of NaR in adipocytes. Although the kidney being the main site for NaR metabolism,^20^ we asked whether adipocytes could take up NaR to generate NAD^⁺^locally. To this end, differentiating 3T3-L1 adipocytes were treated with NaR and harvested at 1, 2, 4, and 6 hours to quantify intracellular NaR and downstream metabolites (**Table S1 and Figure 3A**). There were no detectable endogenous NaR production in adipocytes, while exogenous NaR was rapidly taken up, reaching maximal intracellular levels within one hour. In contrast, intracellular NAD^⁺^ and related metabolites increased more gradually, becoming elevated at 4–6 hours after treatment. These kinetics indicate that adipocytes can take up NaR which subsequently induce NAD^⁺^ metabolism.

**Figure 3.**
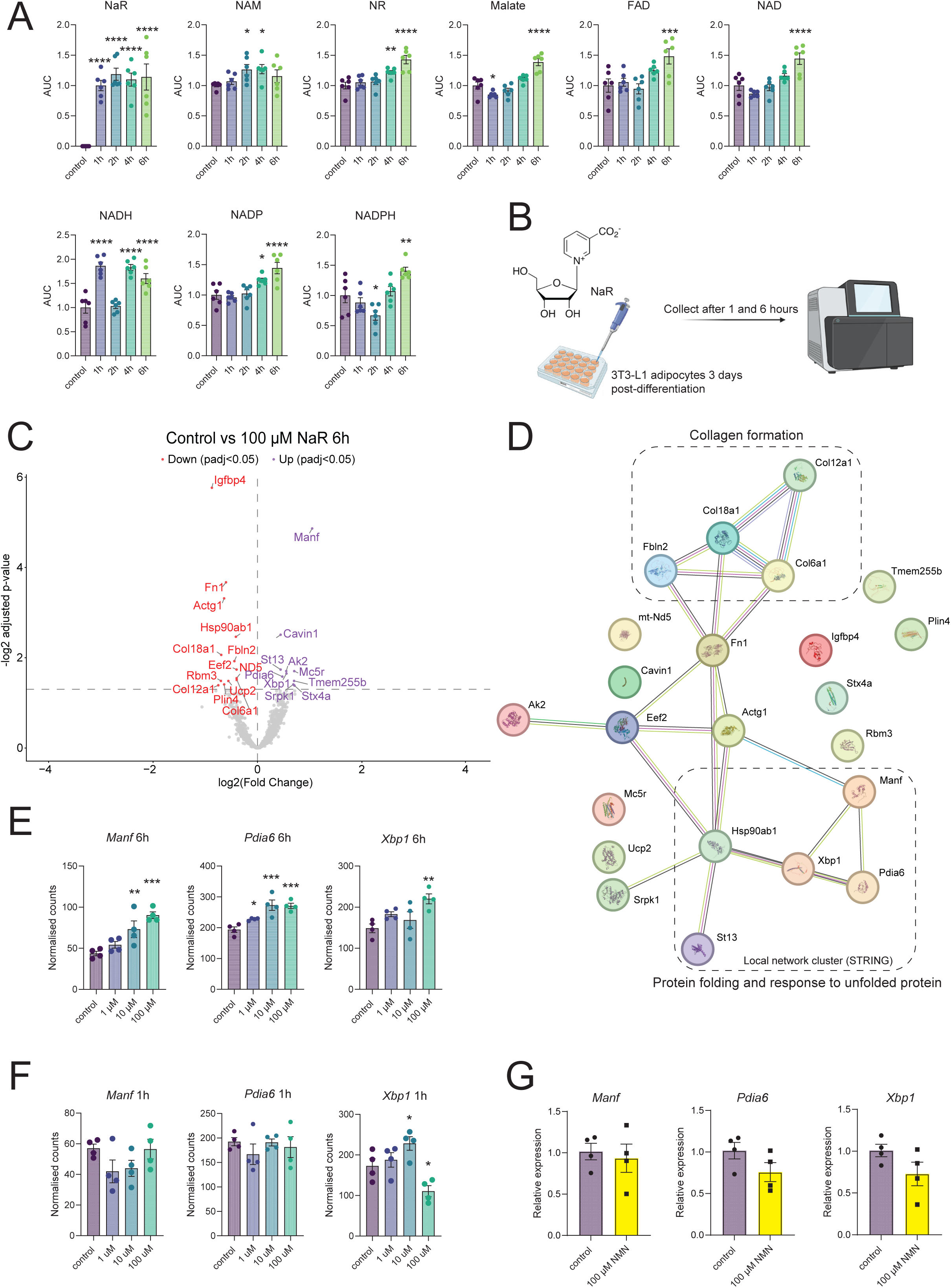
Nicotinic acid riboside (NaR) regulates unfolded protein response (UPR) genes in mouse adipocytes. A) Metabolomic analyses of differentiating 3T3-L1 adipocytes treated with 100 µM NaR for 1, 2, 4 and 6 hours. Six biological replicates per treatment. Statistical comparisons between control and each treatment group were performed using one-way ANOVA with Fisher’s LSD test. *p-value < 0.05, **p-value < 0.01, ***p-value < 0.001, ****p-value < 0.0001. AUC: area under the curve. B) Schematic of the *in vitro* NaR treatment experiment and consequent transcriptomic profiling (Created with BioRender.com). C) Volcano plot displaying differentially expressed genes in 3T3-L1 cells treated with 100 µM NaR for 6 hours compared to their controls. D) Pathway enrichment analysis of the encoded gene products from panel C. E-F) Transcriptomic analyses of *Manf*, *Pdia6* and *Xbp1* from differentiating 3T3-L1 cells treated with NaR at different concentrations for 6 and 1 hours. Four biological replicates per treatment. Statistical comparisons between control and each treatment group were performed using one-way ANOVA with Fisher’s LSD test. *p-value < 0.05, **p-value < 0.01, ***p-value < 0.001. G) Relative expression of UPR genes in differentiating 3T3-L1 cells treated with 100 µM β-NMN for 6 hours. Four biological replicates per group. A student unpaired t-test was performed to determine significant differences between groups.

To define cellular programs responsive to NaR, we performed transcriptomic profiling of differentiating adipocytes treated with increasing concentrations of NaR (0, 1, 10 and 100 µM) and collected after 1 and 6 hours (**Figure 3B and Table S2**). We first focused on the differentially expressed genes in the adipocytes treated with 100 µM for 6 hours compared to control treated cells. Importantly, cell viability was not influenced by this treatment (**Figure S3A**). The analysis revealed a highly selective transcriptional response, with only 23 genes significantly altered (**Figure 3C**). Pathway analysis of the differentially expressed genes and the interaction between their protein products revealed two interconnected clusters, one enriched for collagen formation–associated genes and a second enriched for genes linked to the unfolded protein response (UPR) (**Figure 3D**). Among the UPR-associated genes induced by NaR were *Pdia6* and *Manf*, both of which regulate IRE1α activity and *Xbp1* mRNA splicing^27,28^, as well as *Xbp1* itself. The UPR is a pan-cellular stress response that has been implicated in diverse physiological and pathological contexts, including muscle adaptation to exercise^29^ and hippocampal neurodegeneration.^30^ We therefore measured *Manf*, *Pdia6*, and *Xbp1* expression in myotube (C2C12) and neuronal (HT-22) cell lines treated with NaR (**Figure S3B-D**). No changes were observed in myotubes (**Figure S3C**), whereas neuronal cells showed a trend toward increased *Manf* expression and a significant induction of *Xbp1*, indicating that NaR-mediated regulation of the UPR is not restricted to adipocytes.

In 3T3-L1 cells, expression of *Manf* and *Pdia6* exhibited dose-dependent responses to NaR, while *Xbp1* induction was detectable only at the highest concentration tested (**Figure 3E**), indicating graded engagement of UPR-associated gene programs. Notably, there was no increase in expression at the 1-hour time point and even a decrease in *Xbp1* expression at the highest concentration (**Figure 3F**), suggesting that these effects are either mediated by downstream metabolites or a delayed transcriptional response to NaR. To determine whether induction of these genes reflects a consequence of increased NAD^⁺^ precursor availability, we treated adipocytes with nicotinamide mononucleotide (NMN). In contrast to NaR, 6-hour treatment with NMN did not alter expression of *Manf*, *Pdia6*, or *Xbp1* (**Figure 3G**), indicating that NaR elicits the UPR independent of downstream NAD^⁺^ metabolites.

To assess the *in vivo* relevance of these observations, we examined WAT gene expression in hepatocyte-specific *Bmal1* knockout (LKO) mice and their WT littermates from our previously published dataset.^12^ Under these conditions, *Manf* and *Pdia6* exhibited a distinct peak in expression at ZT16 in WT mice, which was abolished in LKO animals (**Figure S3E**). In contrast, *Xbp1* expression was similar between genotypes. Furthermore, to assess the sufficiency of the hepatocyte clock in regulating WAT UPR expression, we mined our published dataset^12^ consisting of WAT transcriptomics from Bmal1-hepatocyte reconstituted mice (LRE, Bmal1 expression restricted to hepatocytes) and compared them to their WT and full body KO littermates. Hepatocyte-specific *Bmal1* reconstitution was sufficient to restore rhythmic expression of all three transcripts (borderline significance for *Manf*, cosinor P-value 0.058), although with altered phase and amplitude compared to WT, and with *Xbp1* expression remaining more like KOs (**Figure S3F**). Together, these data indicate that hepatic clock function contributes to temporal regulation of UPR-associated genes in WAT, possibly through NaR signaling.

### Proteome Integral Solubility Alteration (PISA) analysis reveals NaR interaction with the prefoldin complex

Our data indicates that NaR induces UPR gene programs through mechanisms that are independent of global changes in NAD^⁺^ metabolism. We therefore asked whether NaR directly engages specific proteins that could account for this transcriptional response. To address this, we performed a proteome integral solubility alteration (PISA) assay to identify NaR-dependent changes in protein thermal stability, a readout used to infer small-molecule–protein interactions.

Differentiating adipocytes were treated with NaR for 1 hour, a time point preceding detectable increases in intracellular NAD^⁺^ levels, followed by protein extraction, thermal challenge, and quantitative proteomics (**Figure 4A**). Across the proteome, 13 proteins displayed increased thermal stability, and 16 proteins were significantly destabilized upon NaR treatment (**Figure 4B and Table S3**). Pathway and protein–protein interaction analysis revealed limited connectivity among the stabilized proteins, which included linker histones such as H1f4 and H1f5, known regulators of chromatin organization (**Figure S4A**). In contrast, the destabilized proteins segregated into two interaction clusters (**Figure 4C**). One cluster comprised extracellular matrix–associated proteins, including Fbn1 and Col1a2. The second cluster contained Pfdn5 and Pfdn6, two core subunits of the prefoldin complex. Prefoldin is a heterohexameric cytosolic chaperone that captures nascent or partially folded polypeptides and delivers them to chaperonins for ATP-dependent folding,^31^ linking it directly to cellular proteostasis. To further assess whether NaR affects prefoldin integrity more broadly, we examined thermal stability changes for individual prefoldin subunits detected in the PISA dataset. All detected prefoldin subunits exhibited reduced stability upon NaR treatment (**Figure 4D and Table S3**), consistent with destabilization of the prefoldin complex as a whole. Such coordinated destabilization could arise from NaR engagement with a specific subunit, resulting in secondary effects on complex assembly or stability. Hence, to determine probability of NaR binding to the prefoldin subunits, we performed docking analyses. We first identified binding pockets where there is a likelihood for a metabolite to bind followed by docking analyses of NaR. NaR was predicted to interact with all prefoldin subunits, including one binding pocket in Pfdn6 (**Figure S4B and Table S4**). To provide additional evidence for the interaction between NaR and individual prefoldin subunits, we performed a modified PISA assay on protein lysates, in which protein complexes are largely disassembled. Lysates were treated with NaR prior to thermal profiling. Under these conditions, only Pfdn1 and 6 displayed significantly increased thermal stability with Pfdn6 being the most significant (**Figure 4E**), suggesting that NaR preferentially associates with Pfdn1 and 6 and that this interaction may underlie the destabilization of the assembled prefoldin complex observed in intact cells.

**Figure 4.**
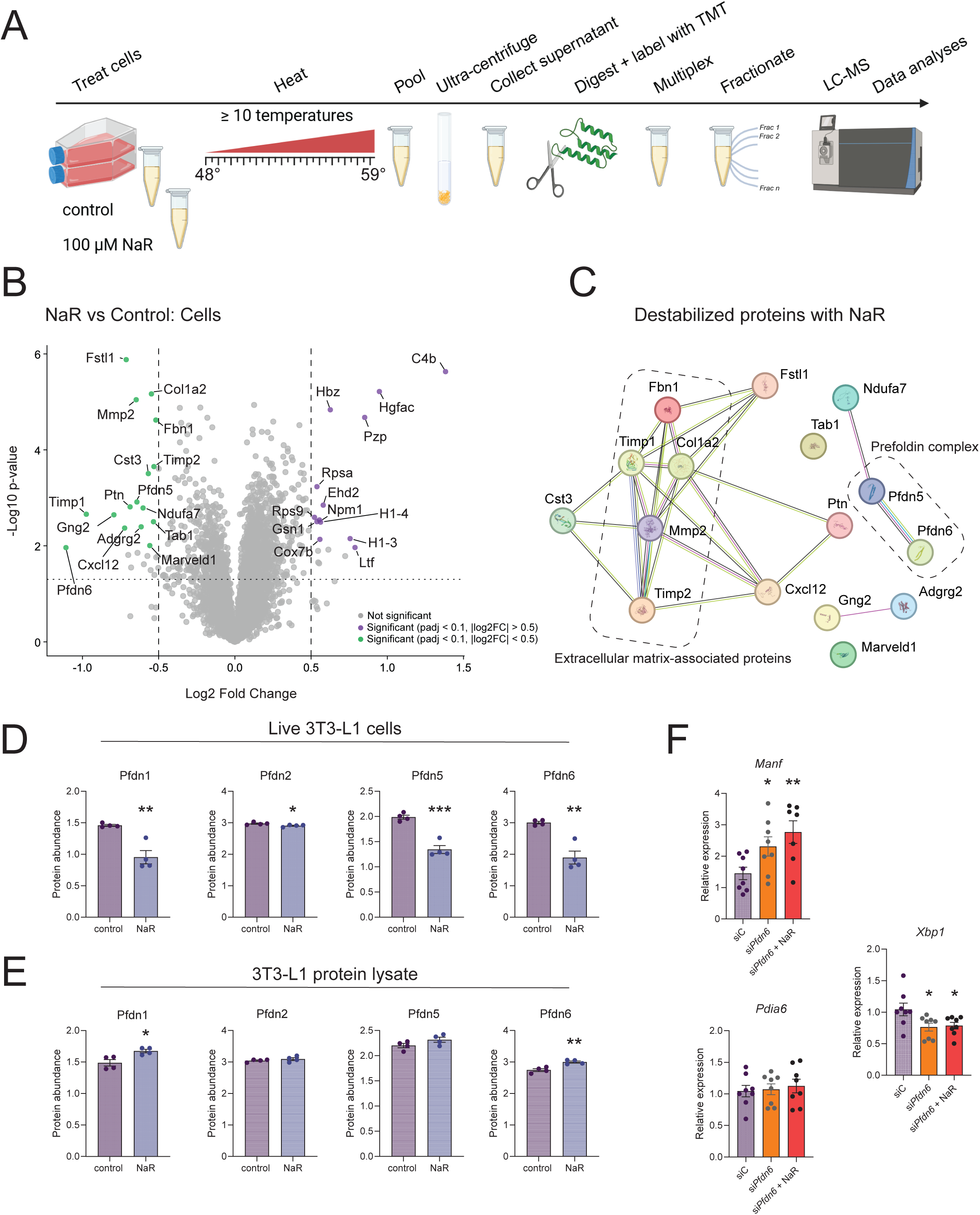
PISA assay demonstrates an interaction between NaR and the prefoldin complex. A) Schematics and workflow of PISA (Created with BioRender.com). B) Volcano plot displaying proteins with differential stability in differentiating 3T3-L1 cells treated with or without 100 µM NaR for 1 hour. C) Pathway enrichment analysis of de-stabilized proteins upon NaR treatment. D) Protein abundance for individual prefoldin subunits detected in the PISA assay performed on living 3T3-L1 cells upon 100 µM NaR treatment. Four biological replicates per group. A student unpaired t-test was performed to determine significant differences between groups. *p-value < 0.05, **p-value < 0.01, ***p-value < 0.001. E) Protein abundance for individual prefoldin subunits detected in the PISA assay performed on cell lysates upon NaR treatment. Four biological replicates per group. A student unpaired t-test was performed to determine significant differences between groups. *p-value < 0.05, **p-value < 0.01. F) Relative expression of UPR genes in differentiating 3T3-L1 control (siC) and *Pfdn6*-KD (si*Pfdn6*) cells. Outliers were identified using the Robust regression and Outlier removal (ROUT). Number of biological replicates for each group is at least seven for each group. Statistical comparisons in relation to siC were performed using one-way ANOVA with Fisher’s LSD test. *p-value < 0.05, **p-value < 0.01.

To determine the involvement of specific prefoldin subunits in regulating the UPR in adipocytes, we performed siRNA-mediated knockdown of Pfdn6 (selecting the most significant subunit in the lysate analysis) in 3T3-L1 preadipocytes, followed by induction of adipogenesis. siRNA treatment resulted in a significant reduction of *Pfdn6* expression (**Figure S4C**), without changes in cell viability compared with non-targeting siRNA (**Figure S4D**) and was associated with induced expression levels of *Manf*, attenuation of *Xbp1* expression, and no changes in *Pdia6* (**Figure 4F**), highlighting the importance of Pfdn6 in modulating the UPR.

### NaR regulates adipocyte Cebpa expression primarily through ATF6

The UPR has been shown to be required for adipogenesis ^32,33^. Although our unbiased RNA-sequencing analysis did not identify broad induction of adipogenic gene programs following NaR treatment (**Figure 3C**), this may reflect the stringent statistical thresholds applied. We therefore revisited the transcriptomic data with a targeted focus on key adipogenic regulators.

Among the adipogenic genes examined (including but not limited to *Pparg*, *Adipoq, Plin1,* and *Fabp4*) NaR stimulation selectively induced expression of *Cebpa*, which we validated by qRT-PCR (**Figure 5A**). This observation is consistent with previous reports demonstrating that XBP1 regulates adipogenesis through *Cebpa*.^32^ Furthermore, CEBPA were one of the destabilized proteins found in our PISA analysis (**Table S3**), suggesting that NaR may contribute to lipid regulation by engaging both the UPR and the adipogenic transcriptional machinery simultaneously. Thus, to determine whether NaR mediates transcriptional induction of *Cebpa* through the prefoldin complex, we combined the *Pfdn6* siRNA treatment with NaR treatment. As hypothesized, in cells treated with non-targeting siRNA, NaR resulted in increased *Cebpa* expression while this effect was abolished in si*Pfdn6* treated cells (**Figure 5B**). We next sought to determine which branch of the UPR mediates this effect. To dissect UPR pathway involvement, we combined NaR stimulation with pharmacological inhibitors targeting the three UPR branches: GSK2656157 (GSK), a PERK inhibitor; Ceapin-A7 (CA7), an inhibitor of ATF6 activation; and 4µ8C, an inhibitor of IRE1 RNase activity (**Figure 5C**). Importantly, there were no differences in viability between the cells treated with either compound (**Figure S4E**). Inhibition of ATF6 with CA7 abolished the NaR induction of all measured UPR genes (**Figure 5D-F**) while inhibition of IRE1 and PERK with 4µ8C and GSK, respectively, did not (**Figure 5D-F**). This suggests that NaR induces UPR through ATF6. However, inhibition of both ATF6 and IRE-1 resulted in sustained suppression of *Cebpa* expression that could not be rescued by NaR treatment (**Figure 5G**). Together, these data indicate that ATF6 and IRE-1 dependent activation is critical for NaR-induced *Cebpa* expression and that this is mediated independent of the UPR response genes. These observations are in line with previous reports highlighting the involvement of these signaling branches in adipogenesis.^32,33^

**Figure 5.**
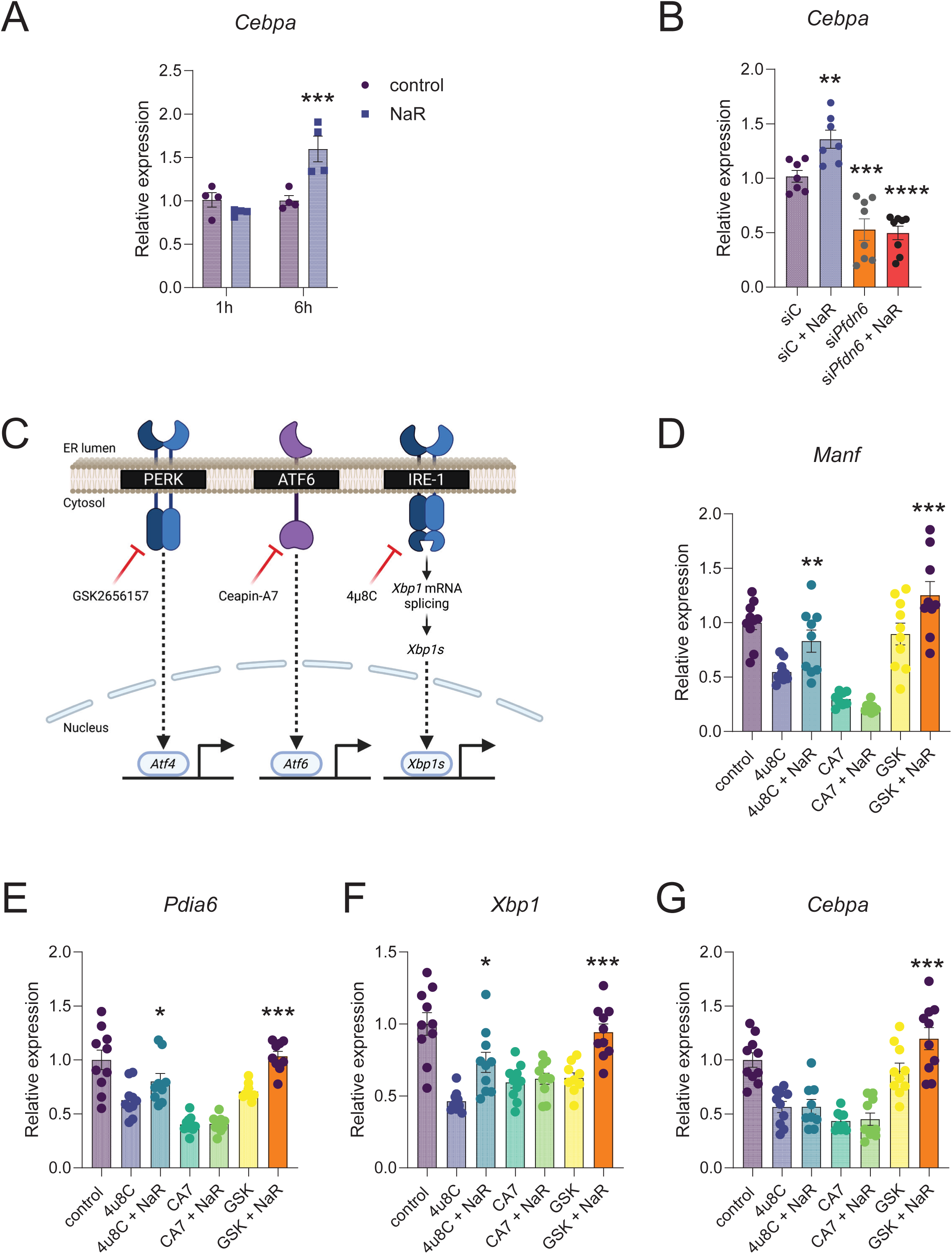
NaR modulates adipocyte *Cebpa* expression through ATF6 signaling. A) Relative gene expression of *Cebpa* measured by qRT-PCR from differentiating 3T3-L1 cells treated with 100 µM NaR for 1 and 6 hours. Four biological replicates per group. Statistical comparisons between the control and each treatment group were performed using two-way multiple-comparison ANOVA with Fisher’s LSD test. ***p-value < 0.001. B) Relative *Cebpa* expression in differentiating 3T3-L1 control (siC) and *Pfdn6*-KD (si*Pfdn6*) cells treated with or without 100 µM NaR for 6 hours. Outliers were identified using the Robust regression and Outlier removal (ROUT). Number of biological replicates for each group is at least seven for each group. Statistical comparisons in relation to siC were performed using one-way ANOVA with Fisher’s LSD test. **p-value < 0.01, ***p-value < 0.001, ****p-value < 0.0001. C) Graphical representation of the ER stress branches and the respective pharmacological inhibitors (in red) targeting them (Created with BioRender.com). D-G) Relative gene expression of *Manf*, *Pdia6*, *Xbp1*, and *Cebpa* in differentiating 3T3-L1 cells treated with UPR inhibitors alone or co-treated with 100 µM NaR for 6 h. Outliers were identified using the robust regression and outlier removal (ROUT) method. Data represent at least eight biological replicates per group. Statistical analyses were performed separately for each gene using two-way ANOVA with treatment (UPR inhibitor) and NaR co-treatment as factors, followed by Fisher’s LSD test to compare each inhibitor with its corresponding co-treated group. *p-value < 0.05, **p-value < 0.01, ***p-value < 0.001.

### Oscillatory NaR signals are integrated by the adipocyte clock to regulate lipid accumulation

The preceding results established that NaR induces UPR-associated gene programs and regulates *Cebpa* expression in adipocytes. We next asked whether the temporal structure of NaR exposure influences adipocyte differentiation outcomes. Given that adaptive versus maladaptive UPR signaling depends on timing and signal resolution,^32–35^ we hypothesized that oscillatory and sustained NaR signals would be interpreted differently by differentiating adipocytes.

To test this, we compared the effects of pulsatile versus constant NaR exposure on lipid accumulation in differentiating adipocytes. 3T3-L1 cells were subjected either to continuous NaR treatment throughout the differentiation period (constant NaR treatment; 8 days) or to a daily 8-hour NaR pulse followed by 16 hours in NaR-free medium (**Figure 6A**). Chronic NaR treatment did not alter CEBPA protein levels in differentiating adipocytes (**Figure 6B**). In contrast, daily pulsatile NaR stimulation resulted in a significant increase in CEBPA protein abundance (**Figure 6C**). Consistent with these effects on CEBPA regulation, quantitative analysis of lipid accumulation revealed reduced lipid area per cell following chronic NaR treatment, whereas pulsatile NaR exposure significantly increased lipid accumulation per cell (**Figure 6D and figures S5A and B**). Together, these data demonstrate that adipocytes differentially interpret NaR signals depending on their temporal structure. Oscillatory NaR exposure promotes adipogenic competence and lipid accumulation, whereas sustained NaR exposure suppresses these processes, underscoring the importance of temporal patterning of systemic metabolic signals in physiology.

**Figure 6.**
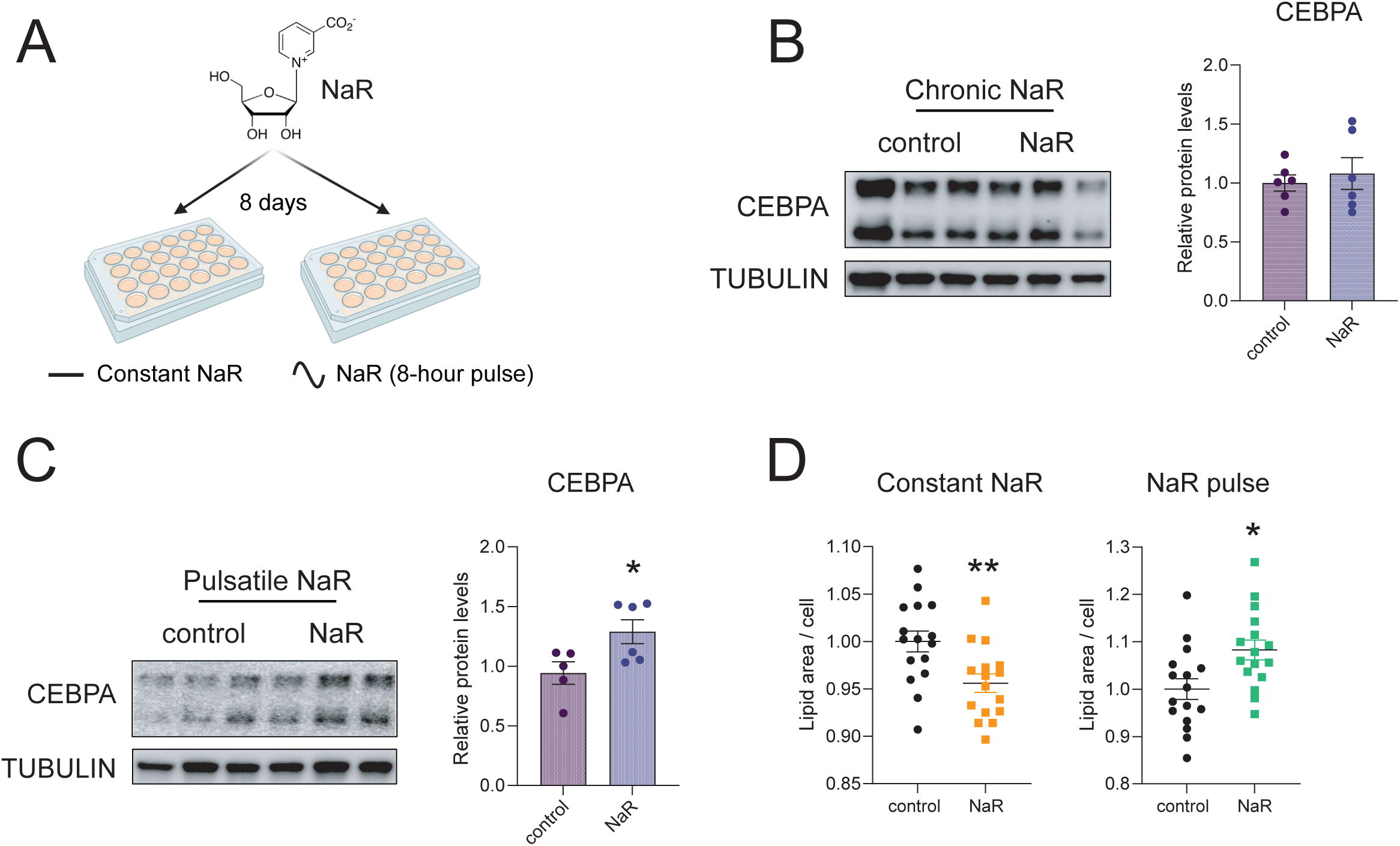
The adipocyte clock integrates oscillatory NaR signaling to regulate lipid accumulation. A) Schematic of the *in vitro* constant and pulsatile NaR protocol (Created with BioRender.com). B-C) Western blot analyses of CEBPA against the reference protein TUBULIN from 3T3-L1 cells subjected to constant or pulsatile NaR protocol for eight days. Outliers were identified using the robust regression and outlier removal (ROUT) method. Data represent at least five biological replicates per group. A student unpaired t-test was performed to determine significant differences between groups. *p-value < 0.01. D) Lipid accumulation analyses of 3T3-L1 cells subjected to constant or pulsatile NaR protocol for eight days. A student unpaired t-test was performed to determine significant differences between groups. *p-value < 0.05, **p-value < 0.01.

## Discussion

Recent advances in circadian physiology highlight that systemic rhythms do not arise solely from isolated tissue clocks but instead emerge from a coupled network involving communication between multiple organs.^14^ While the brain clock is sufficient to drive much of systemic metabolic rhythmicity,^7^ the liver clock has been proposed to function as an integrator of feeding cues that modulates rhythmic outputs in other peripheral tissues.^11^ In support of this model, peptide hepatokines have recently been shown by us and others to contribute to the organization of systemic metabolism.^12,36^ In the present study, we extend the hepatic clock signaling framework by focusing on circulating metabolites as signaling intermediates. Our data identify NaR as a liver clock–controlled metabolite within the hepatic clock–WAT axis and further suggest a mechanistic link to proteostasis through engagement of the prefoldin complex. While previous work has demonstrated a role for the hepatic clock in regulating local UPR signaling within the liver,^37^ our findings raise the possibility that hepatic clock outputs may also influence UPR-associated gene programs in distant tissues. Our results support prior observations, positioning the liver as a central coordinator of systemic circadian rhythms.^4,5,11^

While NaR has recently been shown to circulate and contribute to extrahepatic NAD^⁺^pools,^20^ our data suggest that NaR rhythmicity itself may convey temporal information beyond its role as a metabolic precursor. By demonstrating that NaR oscillations are selectively lost in clock-deficient states and restored by liver clock function, our results raise the possibility that NaR functions as a time-encoded metabolic signal linking hepatic circadian output to systemic regulation of the UPR and adipocyte physiology. In this context, NaR may represent an example of a metabolite whose temporal dynamics, rather than absolute abundance, contribute to inter-organ coordination. An important unresolved question raised by our findings is why the hepatic clock selectively imposes circadian rhythmicity on NaR in circulation, while other NAD^⁺^ intermediates do not exhibit comparable clock dependence. One possibility is that NaR, along with quinolinic acid, serves as a checkpoint for the tryptophan-NAD^⁺^ pathways, buffering metabolic input from the *de novo* NAD^⁺^synthesis pathway. This theory is supported by previous work showing that NaR production is induced in the absence of Nmnat1,^20^ a rate limiting enzyme in the *de novo* NAD^⁺^ synthesis.

Mechanistically, we observed a specific transcriptional response to NaR treatment in adipocytes, inducing the expression of genes involved in the UPR. Furthermore, our PISA analyses indicate that NaR destabilizes the prefoldin complex, yet the functional consequences of this destabilization remain to be fully defined. Prefoldin is traditionally viewed as a constitutive chaperone that delivers nascent or partially folded proteins to downstream folding machinery.^31^ However, destabilization of the prefoldin complex could paradoxically increase its functional flexibility, for example by promoting subunit exchange, transient disassembly, or altered client specificity. In this model, NaR-induced destabilization may render prefoldin more permissive to engage unfolded or aggregation-prone proteins during periods of increased biosynthetic demand, thereby facilitating adaptive proteostasis rather than impairing it. Such a mechanism would be consistent with the selective induction of UPR-associated gene programs observed in response to NaR, without evidence of overt proteotoxic stress. Furthermore, a seemingly counterintuitive observation in our study is that knockdown of *Pfdn6* dampens, rather than enhances, UPR-associated gene expression. Intuitively, loss of a chaperone component might be expected to increase the burden of unfolded proteins and thereby activate stress pathways. One possible explanation is that an intact prefoldin complex is required not only to buffer misfolded proteins but also to generate the specific proteostatic signals necessary to engage adaptive UPR transcriptional programs. In this view, prefoldin may act upstream of UPR signaling as a sensor of folding intermediates, and its loss could blunt the cell’s ability to mount a coordinated transcriptional response. Alternatively, chronic prefoldin deficiency may shift cells into a state of proteostatic adaptation that suppresses inducible UPR signaling, thereby reducing responsiveness to additional inputs such as NaR. Nevertheless, we demonstrate that Pfdn6 plays a crucial role in regulating UPR in adipocytes.

Our observation that NaR induces UPR-associated gene expression in neurons but not in myotubes suggests that NaR acts as a systemic signal whose cellular interpretation is context dependent. While NaR is sufficient to induce *Xbp1* expression in neurons and adipocytes, muscle cells may also require additional cues, namely exercise and the associated engagement of PGC-1α.^29^ Nevertheless, these findings support a model in which NaR serves dual systemic roles: as a metabolic substrate contributing to nicotinamide production in the kidney,^20^ and as a signaling molecule that engages proteostasis pathways in select target tissues.

Our findings underscore the importance of considering time-of-day and temporal signal structure in metabolic research, a principle increasingly recognized across physiological systems.^38^ Classical experimental approaches that rely on static genetic or pharmacological perturbations often overlook the dynamic nature of metabolic regulation and may therefore fail to capture key aspects of physiological or pathophysiological control. In this context, our data provide clarity regarding the dual role of the UPR in adipose biology, reconciling prior observations that UPR signaling can both support and restrain adipocyte differentiation depending on context.^32–35^ Specifically, our results suggest that oscillatory NaR signaling promotes adaptive UPR-associated programs that support adipocyte lipid accumulation, whereas sustained NaR exposure engages a state that is incompatible with efficient lipid deposition. Furthermore, our PISA analysis suggested that NaR interacts with both the prefoldin complex and directly with CEBPA, indicating a coordinated response to NaR in engaging lipid metabolism. Importantly, we propose that these interactions are likely occurring synergistically in response to NaR as *Cebpa* mRNA expression is only induced upon NaR treatment in the presence of the prefoldin subunit Pfdn6. These findings highlight the coordinated temporal dynamics as a critical determinant of metabolic signal interpretation and suggest that therapeutic strategies targeting metabolic pathways may benefit from considering oscillations of molecular processes to better understand homeostatic regulation.^39^

The rhythmicity and adipogenic effects of NaR observed in this study suggest that its physiological role extends beyond NAD^⁺^ biosynthesis to include time-dependent modulation of cellular stress responses and lipid storage. Notably, circulating NaR levels have been reported to decline with age,^20^ raising the possibility that loss of NaR rhythmicity may contribute to age-associated metabolic inflexibility or adipose dysfunction. In this context, alterations in the temporal structure of NaR signaling, rather than changes in absolute metabolite abundance alone, may represent an underappreciated component of metabolic aging. While further work will be required to establish the *in vivo* relevance of these mechanisms, our findings provide mechanistic insights for exploring whether restoration of NaR rhythmicity, or temporal alignment of NaR exposure, can improve metabolic robustness. Future studies aimed at testing time-resolved NaR interventions in models of circadian disruption or aging will be necessary to determine the therapeutic potential of this pathway.

## Limitations

First, although proteome-wide stability profiling (PISA), lysate-based assays, and docking analyses support an interaction between NaR and prefoldin subunits, we have not performed direct biophysical binding measurements to quantitatively define NaR–prefoldin binding affinity. Second, although we show that NaR induces UPR-associated gene expression in multiple cell types, the tissue-specific determinants that shape downstream transcriptional and functional responses require further investigation. Finally, while our in vivo circadian metabolomics establish NaR as a liver clock–controlled circulating metabolite, we have not directly manipulated systemic NaR rhythmicity in vivo to test causal effects on adipose physiology. Future studies employing temporally controlled NaR administration or genetic perturbation of prefoldin in vivo will be necessary to determine the physiological consequences of this signaling axis.

Despite these limitations, our findings establish a mechanistic framework linking systemic circadian signaling to metabolite-dependent regulation of cellular homeostasis, highlighting the interactions between oscillatory and linear molecular processes.

## Materials and methods

### Cell studies

#### Culture and differentiation of the murine 3T3-L1 adipocyte cell line

3T3-L1 cells were cultured as previously described.^12^ Cells were grown in growth media consisting of DMEM/F12, 10% donor calf serum, 2 mM l-glutamine, 1X Penicillin-Streptomycin Solution (Gibco) and 3151 mg/L D-Glucose. Once cells reached 90% confluence, they were washed with 1X phosphate-buffered saline (PBS) and exposed to differentiation medium consisting of DMEM/F12, 10% fetal bovine serum, 3151 mg/L D-Glucose, 100 μM IBMX (Sigma Cat #7018), 1 μM dexamethasone (Sigma Cat #D1756), 1.75 nM bovine insulin (Sigma Cat # I6634) and 10 µM rosiglitazone (Cayman Chemicals Comp. USA, #71740). Three days after initiating differentiation, the media was removed and was replaced with differentiation medium with 1.75 nM insulin for two more days. Media was replaced on every other day basis.

#### Culture of the murine C2C12 muscle cell line

As previously described,^40^ C2C12 myoblasts were cultivated in Dulbecco’s modified Eagle’s medium (Thermo-Fisher Scientific) supplemented with 10% fetal bovine serum and containing 4.5 g/L glucose, 4 mM l-glutamine, 1 mM sodium pyruvate and 1% antibiotic-antimycotic solution (Gibco). Myogenic differentiation was induced in fully confluent cells by switching the fetal bovine serum in the growth medium for 2% horse serum. The NaR treatment experiments were performed after 5 days of differentiation.

#### Culture of the murine HT-22 hippocampal neuronal cell line

HT-22 cells were cultured, according to the suppliers protocol (Millipore Sigma), in high glucose Dulbecco’s modified Eagle’s medium (Sigma), 10% fetal bovine serum, 1X L-Glutamine and 1X Penicillin-Streptomycin Solution (Gibco). Cells were cultivated on 24-well culture plates. After 90% confluence was reached, cells were treated with 100 µM NaR for 6h and further processed for RNA extraction.

### Cell treatments

3T3-L1 cells were treated with 1, 10 and 100 µM NaR (Toronto Research Chemicals Inc., Canada, Cat #TRC-N429400) as well as with β-nicotinamide mononucleotide (NMN) (Sigma Cat #N3501) for 1h and 6h three days post differentiation initiation. Controls cells were treated with differentiation media and water as both NaR and β-NMN were diluted in DNase/RNase free water. Subsequently, cells were lysed with lysis buffer containing 2-mercaptoethanol and stored immediately at −20 until further processing.

To assess the continuous effect of NaR on lipid accumulation, cells were differentiated with 100 µM NAR for 8 days. As for the pulse protocol, cells were only exposed to NaR for 8 hours daily, after which cells were washed with differentiation medium and exposed to a fresh one without NaR. Control cells were treated the same but without being exposed to NaR. All experiments were carried out 8 days post differentiation induction.

Incubations with inhibitors of the UPR (4μ8C [64 μmol/l], ceapin-A7 [4.7 μmol/l], GSK2656157 [5 μmol/l]) were performed for 6h three days post differentiation initiation. Inhibitor concentrations were selected based on results from published studies.^41^

### Reverse transfection of 3T3-L1 murine adipocytes

Reverse transfection of 3T3-L1 pre-adipocytes was performed one day prior to beginning of differentiation. Transfection media was prepared using Pure DMEM Medium, siRNA (non-targeting control and PFDN6 40 nM final concentration) and Dharmafect 1 (Horizon Discovery, US). Thereafter, the mixture was plated and incubated at room temperature for 30 min before adding cells resuspended in proliferation media at a concentration of 250 cells/μL. After 24 h, cells were washed once with 1x PBS and differentiation cocktail added on cells. At day 3, cells were treated with NaR for 6 hours and further processed for RNA analyses.

### RNA isolation, cDNA synthesis and real-time qPCR

RNA from cells was extracted using commercial kits (MACHEREY-NAGEL GmbH & Co. KG) according to the manufacturer’s instructions. The concentration, quality and purity were measured using Nanodrop 2000 (Thermo Fisher Scientific, Lafayette, CO). 300 ng of RNA isolated from cells were reverse transcribed with iScript cDNA synthesis kits (Bio-Rad, Hercules, CA). Quantitative real-time polymerase chain reaction (qRT-PCR) analysis was performed using CFX384 Touch Real-Time PCR Detection System (Bio-Rad, Hercules, CA) with SYBR Green Master Mix (Applied Biosystems). The list of used qRT-PCR primers is: *Cebpa* (Fwd – CAAGAACAGCAACGAGTACCG, Rev –GTCACTGGTCAACTCCAGCAC), *Xbp1* – (Fwd – AGCAGCAAGTGGTGGATTTG, Rev – GAGTTTTCTCCCGTAAAAGCTGA), *Manf* (Fwd – TCTGGGACGATTTTACCAGGA, Rev – TCTTGCTTCACGGCAAAACTTTA), *Pdia6* (Fwd – AGCTGCACCTTCTTTCTAGCA, Rev – CAGGCCGTCACTCTGAATAAC), *Pfdn6* (Fwd – CTATGTCAGGGAGGCAGAAGCT, Rev –CAGCTCCTGTTTGACAAGCACG) and normalized against the housekeeping genes *18s* (Fwd – CGCCGCTAGAGGTGAAATTC, Rev – CGAACCTCCGACTTTCGTTCT), *Lrp10* (Fwd – GGATCACTTTCCCACGTTCTG, Rev – GAGTGCAGGATTAAATGCTCTGA) and *Hprt* (Fwd – TCAGTCAACGGGGGACATAAA, Rev – GGGGCTGTACTGCTTAACCAG).

### Protein extraction, quantification, gel electrophoresis and western blotting

Cells plated in 6-well plates were lysed on ice using RIPA buffer supplemented with Protease Inhibitor Cocktail (Roche) and phosphatase inhibitors (Millipore Sigma). Protein concentration was estimated using the Pierce BCA Protein Assay Kit (Thermo Fisher Scientific, USA). Samples were diluted to normalize volume and concentration, followed by denaturation at 95 °C for 5 min with the addition of 4X Laemmli and 2-mercaptoethanol. Proteins were separated by SDS–PAGE electrophoresis and transferred to nitrocellulose membranes (Bio-Rad, Hercules, CA). Membranes were incubated in blocking reagent (3% of Amersham ECL Prime Blocking solution reagent in Tris-buffered saline-tween 20 (TBS-tween)) for 1 h, then in primary antibody overnight at 4°C. The antibodies used were anti-CEBPA, Cell Signalling, #8178 and anti-alpha-TUBULIN, Cell Signalling, #9099. After three washes in TBS-tween (10 min each), membranes were incubated in anti-rabbit IgG, HRP linked, Cell Signaling Technologies, 7074S for 1 h at room temperature in 3% blocking solution. Membranes were incubated with ECL western-blotting substrate (Amersham International, GE Healthcare) and images were acquired using a ChemiDoc MP Imaging System (Bio-Rad Laboratories).

### Assessment of lipid area

3T3-L1 pre-adipocytes were plated in Corning® CellBIND® 96 well plates until 90% confluence was reached. Thereafter, cells were either exposed to a continuous or 8-hour pulsatile NaR treatment protocol for 8 days. Cells were then fixed on the plates using 4% paraformaldehyde for 15 minutes and stained with Hoechst and BODIPY for 20 minutes, then preserved in PBS. High – throughput quantification of nuclear (Hoechst) and lipids (BODIPY) staining were carried out using a CellInsightCX5 High Content Screening Platform at 10X magnification.

### Cell viability assay

3T3-L1 adipocytes were seeded in a 96-well plate at 0.5 x 10^4^ per well. Three days post differentiation induction, cells were treated with different compounds for 4 hours. Cell viability was evaluated using Cell Counting Kit-8 (HY-K0301, MedChemExpress, USA) according to the manufacturer’s instructions. Briefly, 10μL of CCK-8 solution was added to culture medium and incubated for 2 h. The absorbance at 450 nm wavelength was determined with a reference wavelength of 570 nm. Each well represented an independent biological replicate. Absorbance values were blank-corrected and normalised to the mean of untreated control wells from the same experiment, which were set to 100% viability.

### Targeted metabolomic analyses

3T3-L1 cells, 3 days post differentiation induction, were treated with 100 µM NaR for 1, 2, 4 and 6 hours and were lysed with ice-cold methanol and immediately snap frozen in liquid nitrogen. The tubes were stored at −80 °C until their analysis. On the day of analysis, the tubes were vortexed for 5 s and subsequently sonicated in an ice-cold water ultrasound bath. The samples were then vortexed for an additional 5 s and centrifuged at 12,000 × g for 15 min at 8 °C. Finally, 200 μL of the supernatant was transferred into amber LC–MS vials equipped with 300 μL inserts for subsequent analysis. Table 1 provides a summary of the compounds detected, where the values are provided as area under the curve in arbitrary units (A.U.) for the polar compounds and concentrations in nanogram per pellet for NaR.

### Proteome Integral Solubility Alteration (PISA) assay

The PISA assay was performed as previously described.^42,43^Briefly, 3T3-L1 cells were grown and differentiated in a T175 cell culture flask (Sigma). At day 3 of their differentiation stage, they were treated with 100 µM NaR for 1 hour. Afterwards, cells were washed with warm 1X PBS and resuspended in PBS. The cell suspensions were freeze-thawed in liquid nitrogen 5 times and then centrifuged at 10,000 g for 5 min to remove the cell debris. The protein concentration in the lysate was measured using Pierce BCA assay (Thermo Fischer). The cleared lysate was then aliquoted in 3 replicates and treated with the NaR for 30 min at 37°C in 350 μL reaction volume. After the reaction, the samples from each replicate were aliquoted into 10 wells in a 96-well plate and heated for 3 min in an Eppendorf gradient thermocycler (Eppendorf; Mastercycler X50s) in the temperature range of 48-59°C. Samples were then cooled for 3 min at RT and afterwards snap frozen and kept on ice. Samples from each replicate were then combined and transferred into polycarbonate thickwall tubes and centrifuged for 20 min at 100,000 g and 4°C.

The soluble protein fraction was transferred to new Eppendorf tubes. Protein concentration was measured in all samples using Pierce BCA Protein Assay Kit (Thermo), the volume corresponding to 30 µg of protein was transferred from each sample to new tubes and urea was added to a final concentration of 4 M. Dithiothreitol (DTT) was added to a final concentration of 10 mM and samples were incubated for 1 h at RT. Subsequently, iodoacetamide (IAA) was added to a final concentration of 50 mM and samples were incubated at RT for 1 h in the dark. The reaction was quenched by adding an additional 10 mM of DTT. Proteins were precipitated using methanol/chloroform and resuspended in 20 mM EPPS containing 8M urea. Subsequently, urea was diluted to 4M using EPPS and Lysyl endopeptidase (LysC; Wako) was added at a 1:75 w/w ratio and samples were incubated at RT overnight. Samples were diluted with EPPS to the final urea concentration of 1 M, and trypsin was added at a 1:75 w/w ratio, followed by incubation for 6 h at RT. Acetonitrile (ACN) was added to a final concentration of 20% and TMT reagents were added 4x by weight (120 μg) to each sample, followed by incubation for 2 h at RT. The reaction was quenched by the addition of 0.5% hydroxylamine. Samples were combined, acidified by TFA, cleaned using Sep-Pak cartridges (Waters) and dried using DNA 120 SpeedVac Concentrator (Thermo).

For PISA assay in living cells, cells were cultured in 6-well plates to a density of 250,000 cells per plate. At day 3 of their differentiation stage, they were treated with 100 µM NaR for 1 hour. Cells were then washed with PBS, scraped off and resuspended in PBS. The samples were divided into 10 aliquots in PCR plates and heated like above, then snap-frozen and kept on ice. The samples from each replicate were then pooled and 0.4% final concentration of NP40 was added, followed by three further freeze-thaw cycles. The rest of the protocol was identical to PISA in lysate.

### LC-MS/MS analysis and data acquisition

TMT-labeled peptides were fractionated by high-pH reversed phase separation using a XBridge Peptide BEH C18 column (3.5 um, 130 Å, 1 mm x 150 mm, Waters) on an Ultimate 3000 LC system (Thermo Scientific) at a flow of 42 ul/min. Peptides were separated using the following gradient: 2% B to 15% B over 3 min to 45% B over 59 min to 80% B over 3 min followed by 9 min at 80% B then back to 2% B over 1 min followed by 15 min at 2% B. Buffer A was 20 mM ammonium formate in water, pH 10 and buffer B was 20 mM ammonium formate in 90% acetonitrile, pH 10. A total of 72 fractions were collected, pooled into 24 fractions using a post-concatenation strategy as previously described (Wang et al., 2011) and dried under vacuum.

Dried peptides were resuspended in 0.1% formic acid in water and subjected to LC–MS/MS analysis using an Orbitrap Eclipse Tribrid Mass Spectrometer fitted with a Vanquish Neo system (both Thermo Scientific) and a custom-made column heater set to 60°C. Peptides were resolved on a homemade RP-HPLC column (75 um × 30 cm) packed with C18 resin (ReproSil Saphir 100 C18, 1.5 um, Dr. Maisch GmbH) at a flow rate of 0.2 ul/min. Peptides were separated using the following gradient: 2% B to 8% B over 5 min to 25% B over 70 min to 35% B over 15 min to 95% B over 1 min followed by 4.5 min at 95% B then back to 2% B over 1 min. Buffer A was 0.1% formic acid in water and buffer B was 80% acetonitrile, 0.1% formic acid in water.

The mass spectrometer was operated in DDA mode with a cycle time of 3 s between master scans. MS1 spectra were acquired in the Orbitrap at a resolution of 120,000, a scan range of 400 to 1600 m/z, AGC target set to “Standard” and maximum injection time set to “Auto”. Precursors were filtered with precursor selection range set to 400–1600 m/z, monoisotopic peak determination set to “Peptide”, isolation window center set to “Most Abundant Peak”, charge state set to 2 - 6, a dynamic exclusion of 60 s, a precursor fit of 70% in a window of 0.7 m/z and an intensity threshold of 2.5e4. Precursors selected for MS2 analysis were isolated in the quadrupole using a 0.7 m/z isolation window and fragmented using HCD with a normalized collision energy of 35%. MS2 spectra were acquired in the Orbitrap in centroid mode at a resolution of 30,000 with scan range mode set to “Define First Mass”, a first mass of 110 m/z, normalized AGC target set to 200% and a maximum injection time of 54 ms.

The acquired raw files were analysed using the SpectroMine software (Biognosis AG, Schlieren, Switzerland). Spectra were searched against a murine database consisting of 17085 protein sequences (downloaded from Uniprot on 20220222) and 392 common contaminants. Standard Pulsar search settings for TMT 16 pro (“TMTpro_Quantification”) were used and resulting identifications and corresponding quantitative values were exported on the PSM level using the “Export Report” function. Acquired reporter ion intensities were employed for automated quantification and statistical analysis using the in-house developed SafeQuant R script (v2.3) (Ahrné et al., 2016). This analysis included adjustment of reporter ion intensities, global data normalization by equalizing the total reporter ion intensity across all channels, data imputation using the knn algorithm, summation of reporter ion intensities per protein and channel and calculation of protein abundance ratios. To meet additional assumptions (normality and homoscedasticity) underlying the use of linear regression models and t-tests, MS-intensity signals were transformed from the linear to the log-scale. Protein intensities were normalized based on the total intensity of each sample and then fold changes and adjusted p values were calculated by log2 transformation and median centering of protein intensities followed by students t-test and Benjamini-Hochberg adjustments.

### Pfdn6 structure preparation, pocket prediction and molecular docking

The predicted structure of Pfdn6 was obtained from AlphaFold. Putative ligand-binding pockets were identified using PrankWeb,^44^ which predicts binding sites based on structural and physicochemical features of the protein. The highest-ranked pocket was selected for further analysis.

Molecular docking was performed using SwissDock^46,48^ to evaluate the potential interaction between prefoldin proteins and NaR. The AlphaFold-predicted structure of Pfdn1, 2, 4, 5 and 6 was used as the receptor, and the SMILES representation of NaR ([O-]C(C1=C[N+]([C@@H]2O[C@@H]([C@H]([C@H]2O)O)CO)=CC=C1)=O) was used as the ligand. Docking was performed using default parameters, and predicted binding modes were analyzed based on estimated binding free energy and pose stability.

### RNA sequencing

RNA samples from 3T3-L1 cells treated with 1, 10 and 100 µM NaR for 1h and 6h were sequenced using the Smart-seq3 method.^45^ Raw Fastq files were processed with zUMIs (v2.4.1 or newer) and STAR (v2.5.4b) to generate expression profiles. UMI-containing reads were identified using zUMIs with specified patterns and definitions. Mouse cDNA was mapped against the mm10 genome with CAST SNPs masked to avoid bias. Data were quantified using Ensembl GRCm38.91 annotations.

### Rhythmicity and Deseq2 analyses

The dryR package^47^ was used to identify differential rhythmicity in serum and liver metabolomics data collected in a time series manner using raw count data, with default parameters. The dryR function implements a rhythmicity analysis based on a generalized linear model with a subsequent model selection using the Bayesian information criterion. The raw normalised data from Metabolon for serum^7^ and liver^15^ samples was used to identify rhythmic metabolites with the dryR package.

To detect differentially expressed genes in the generated transcriptomic data from 3T3-L1 cells treated with 1, 10 and 100 µM NaR, the Smart-seq3 count data was analysed using the DESeq2 package.^49^

### Statistical analyses

All data are displayed as mean ± S.E.M. For each experiment, the number of biological replicates, statistical test and significance threshold can be found in the figure legends. Unless otherwise stated, data were analysed in Prism 10.1 (GraphPad) and R Studio.

### Data availability

RNA-seq data will be deposited to Gene Expression Omnibus (GEO) before publication. Proteomic data have been deposited to the ProteomeXchange Consortium (https://www.proteomexchange.org/) via the MassIVE partner repository (https://massive.ucsd.edu/) with MassIVE data set identifier MSV000100889 and ProteomeXchange identifier PXD074620.

## Acknowledgements

We would like to thank the Single cell Sequencing Core facility of Flemingsberg campus (SICOF), Karolinska Institutet for their services. This facility is supported by the Karolinska Institutet Infrastructure council. We would also like to acknowledge the contributions of the Small Molecule Mass Spectrometry Core Facility (KI-SMMS), financed by the Infrastructure Board at Karolinska Institutet for providing scientific input and for the metabolomics analysis of the samples. We would like to thank the Proteomics Core Facility at the Biozentrum, University of Basel, Switzerland for the collaborative analysis of the proteomics samples. Further, we thank Professor Anna Krook and Dr. Nicholas Pillon for kindly providing us with 3T3-L1 and C2C12 cell lines. Finally, we would like to acknowledge the financial support provided to Dr. Paul Petrus by the Novo Nordisk foundation (NNF22OC0073101), Karolinska Institutet, The Swedish Research Council (202302700) The Wenner-Gren foundations (FT2022-0001), the Åke Wiberg foundation and Jeanssons Stiftelser (J2023-0136). A.A.S. acknowledges funding from the Novo Nordisk foundation (NNF25OC0100921) and the Swedish Research Council (2023-02692).

## Author contribution

P.P. and I.V. conceived the project and wrote the original draft of the manuscript. All authors reviewed and approved the final version. I.V., C.S., L.Z., A.A.S., and D.R. performed experiments and data analyses. P.P., A.A.S., and A.S. secured funding for the study.

---

**Figure S1.**
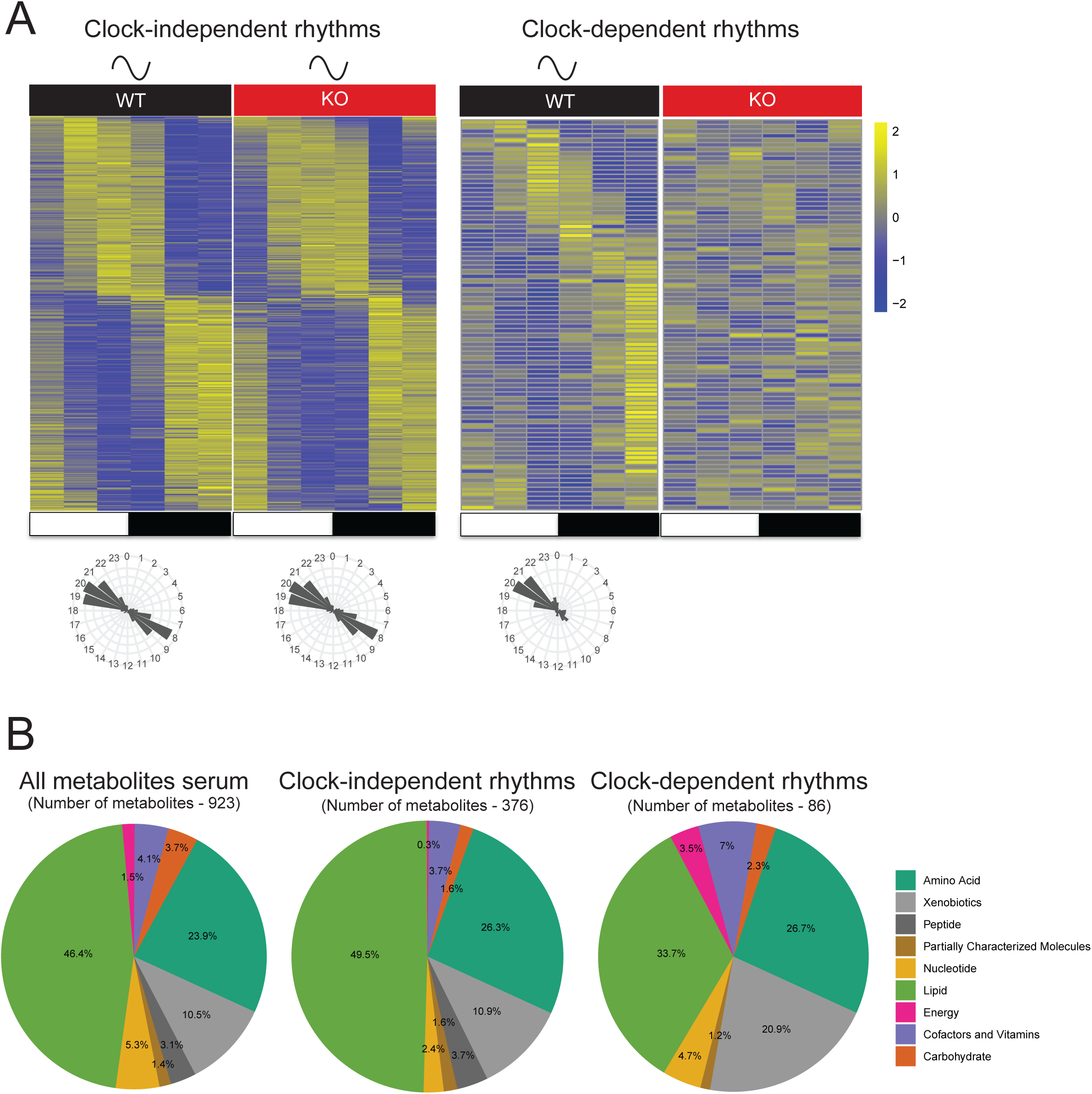
Post-hoc circadian analyses reveal distinct feeding-driven and clock-dependent metabolic oscillations. A) Phase-sorted heatmaps of serum metabolic rhythms in both genotypes (clock-independent) or WT only (clock-dependent). The radar charts illustrate the number of rhythmic metabolites with different peak phases within the groups. B) Pie charts depicting the distribution of metabolite classes from panel A among the clock-independent and clock-dependent metabolites, relative to the total detected metabolite pool.

**Figure S2.**
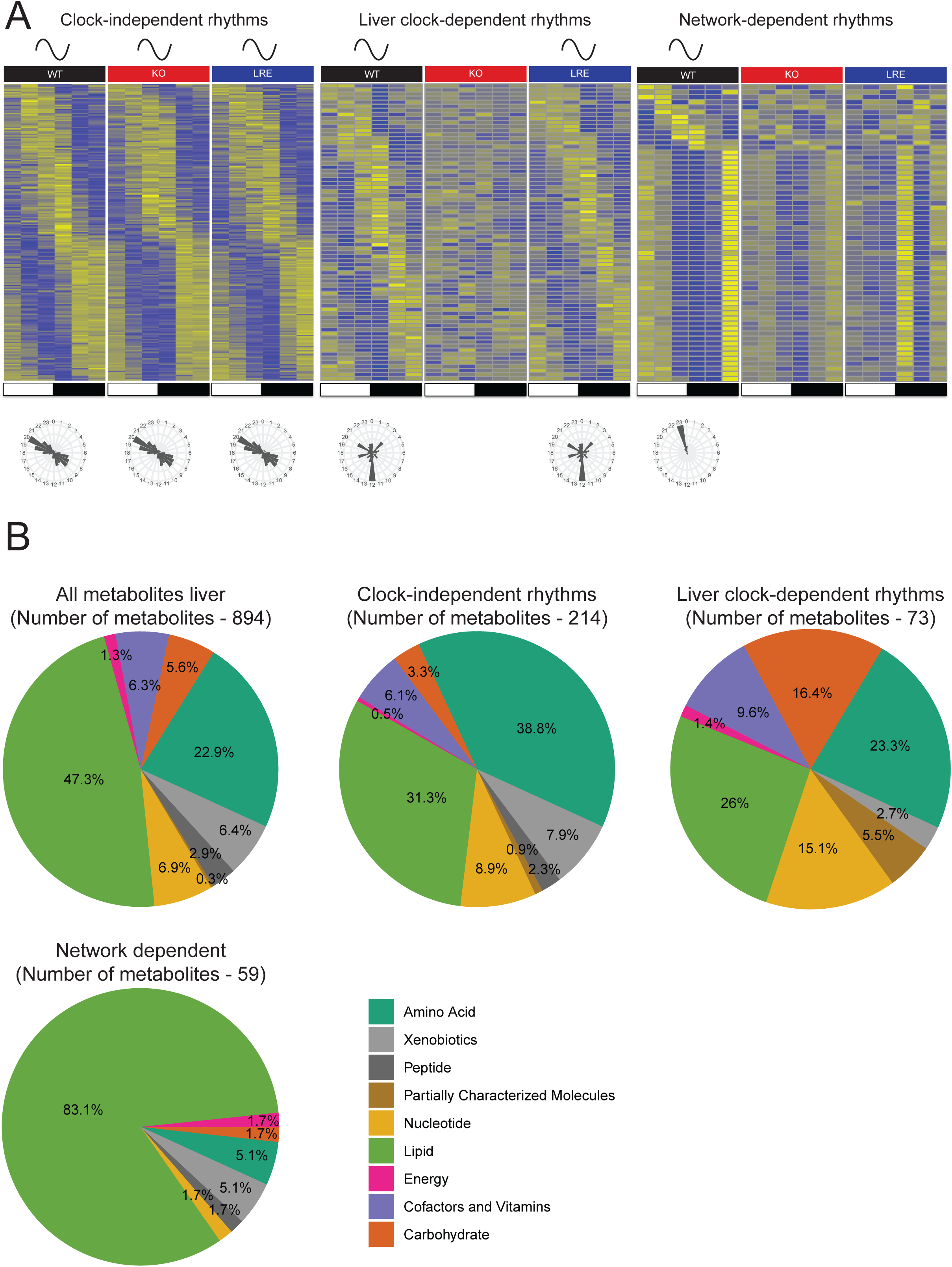
Post-hoc circadian analyses reveal distinct feeding-driven and liver clock-dependent metabolic oscillations. A) Phase-sorted heatmaps and corresponding radar plots of rhythmic liver metabolites categorized as: clock-independent (rhythmic across all genotypes), liver clock-dependent (shared by WT and LRE), and network-dependent (WT-specific). The radar charts illustrate the number of rhythmic transcripts with different peak phases within the groups. B) Pie charts representing the distribution of metabolite classes from panel A among clock-independent, liver clock-dependent and network dependent metabolites, relative to the total detected metabolite pool.

**Figure S3.**
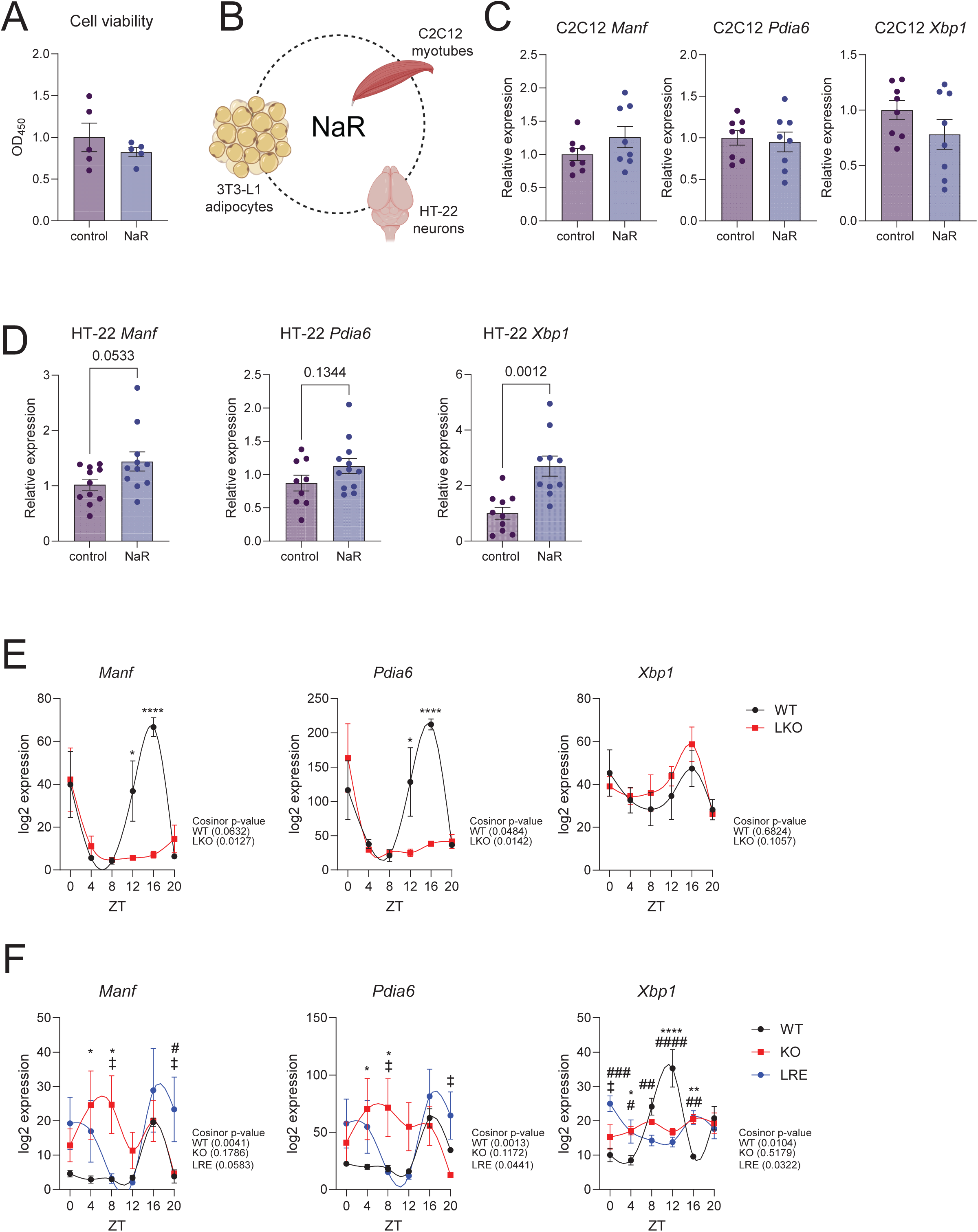
Systemic NaR regulation of the UPR across tissues. A) Cell viability of differentiating 3T3-L1 cells treated with 100 µM NaR for 6 hours. Five biological replicates per group. A student unpaired t-test was performed to determine significant differences between groups. B) Schematic of the *in vitro* approach (Created with BioRender.com). C) qPCR gene expression of UPR genes in differentiated murine C2C12 myotubes. A student unpaired t-test was performed to determine significant differences between groups. D) qPCR relative expression of UPR genes in murine HT-22 hippocampal cells. A student unpaired t-test was performed to determine significant differences between groups. P-values are indicated in each individual graph. E) WAT log2 expression of *Manf*, *Pdia6* and *Xbp1* in WT (black) and LKO (red) mice. Outliers were identified using the Robust regression and Outlier removal (ROUT). Number of biological replicates for each group is at least four for each timepoint. Graphs represent mean log2 expression values and error bars represent standard error of the mean. Cosinor *p*-values are shown; statistical comparisons between groups at individual time points were performed using two-way ANOVA with Fisher’s LSD test. *Represents significant differences in the comparison between WT and KO. *p-value < 0.05, ****p-value < 0.0001. F) WAT log2 expression of *Manf*, *Pdia6* and *Xbp1* in WT (black), KO (red) and LRE (blue) mice. Outliers were identified using the Robust regression and Outlier removal (ROUT). Number of biological replicates for each group is at least three for each timepoint. Graphs represent mean log2 expression values and error bars represent standard error of the mean. Cosinor *p*-values are shown; statistical comparisons between groups at individual time points were performed using two-way ANOVA with Fisher’s LSD test. *Represents significant differences in the comparison between WT and KO, ‡ represents significance in the comparison between KO and LRE and # represents significance in the comparison between WT and LRE. *p-values < 0.05, **p-value < 0.01, ****p-value <0.0001, ^‡^p-value < 0.05, ^#^p-value, ^##^p-value < 0.01, ^###^p-value < 0.001, ^####^p-value <0.0001.

**Figure S4.**
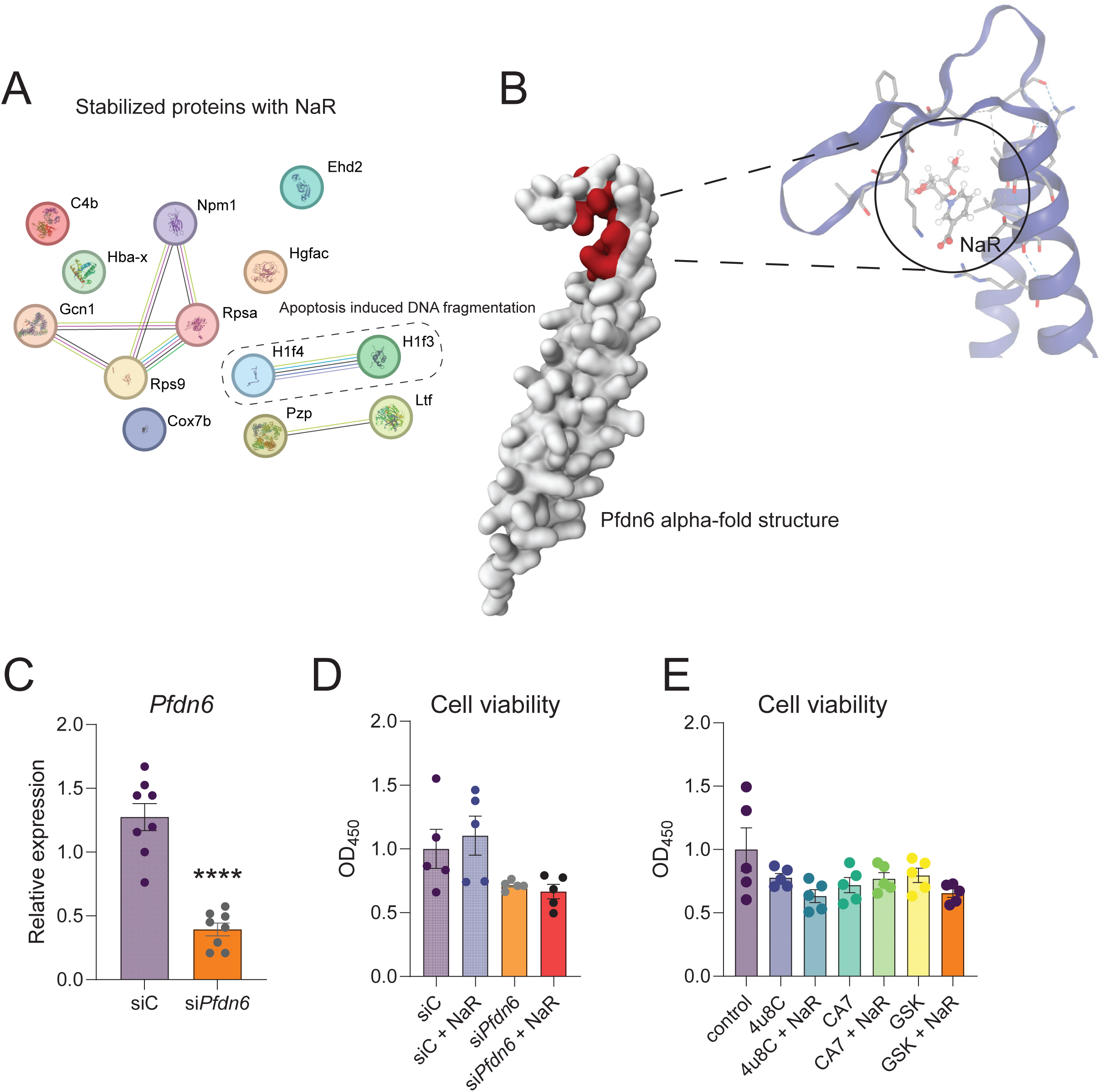
PISA reveals an interaction between NaR and the prefoldin complex. A) Pathway enrichment analysis of stabilized proteins upon NaR treatment. B) The AlphaFold-predicted structure of Pfdn6 is shown with the highest-ranked ligand-binding pocket identified by PrankWeb highlighted in red. SwissDock AutoVina docking analysis demonstrates that NaR can be spatially accommodated within the predicted pocket. C) Relative expression of *Pfdn6* in differentiating 3T3-L1 control (siC) and *Pfdn6*-KD (si*Pfdn6*) cells. Eight biological replicates per group. A student unpaired t-test was performed to determine significant differences between groups. ****p-value < 0.01. D) Cell viability of differentiating 3T3-L1 control (siC) and *Pfdn6*-KD (si*Pfdn6*) cells treated with 100 µM NaR for 6 hours. Five biological replicates per group. Statistical comparisons in relation to siC were performed using one-way ANOVA with Fisher’s LSD test. E) Cell viability of differentiating 3T3-L1 cells treated with UPR inhibitors alone or co-treated with 100 µM NaR for 6 h. Five biological replicates per group. Statistical analyses were performed using two-way ANOVA with treatment (UPR inhibitor) and NaR co-treatment as factors, followed by Fisher’s LSD post hoc test to compare each inhibitor with its corresponding co-treated group.

**Figure S5.**
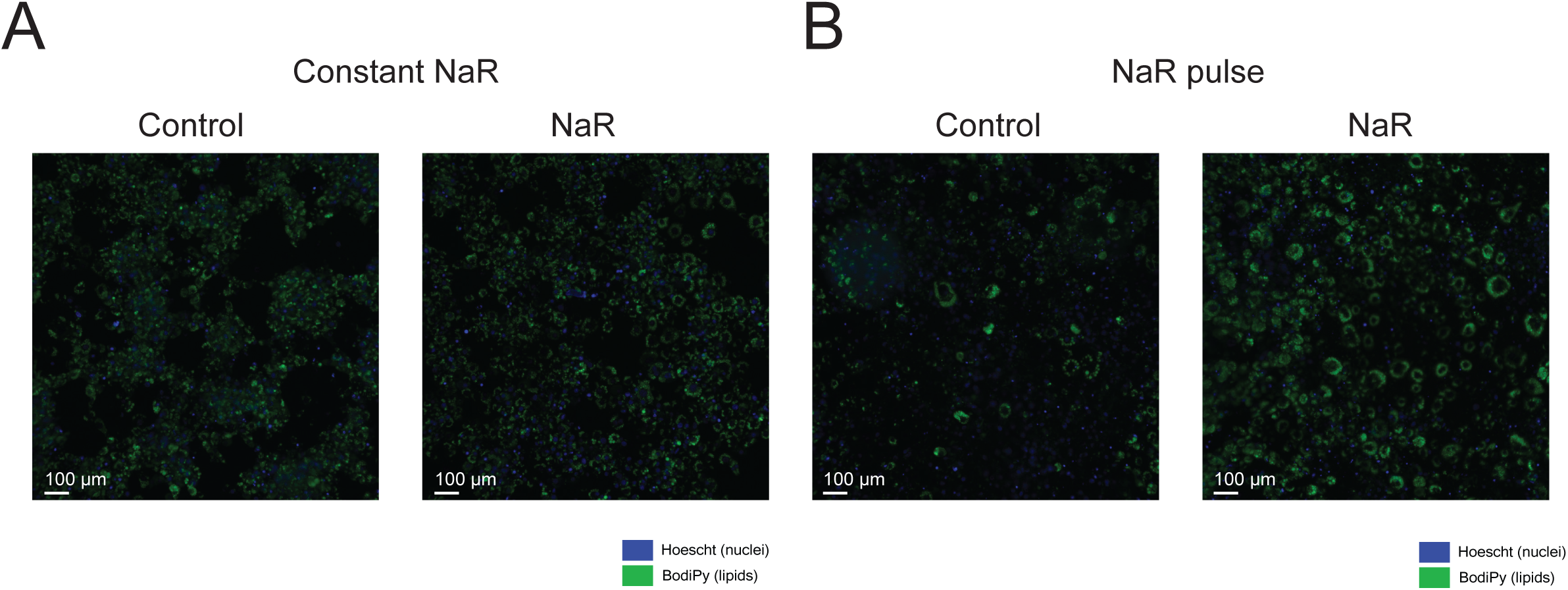
Oscillatory NaR signals regulate lipid accumulation. A-B) Images of 3T3-L1 cells at 10X magnification. Hoescht (Nuclei, blue) and Bodipy (lipid droplet, green) staining of differentiated 3T3-L1 adipocytes subjected to constant or pulsatile NaR protocol for eight days, throughout the differentiation period. The scale bars on all images are 100 μm.

